# Whole mitochondrial genome sequencing identifies unique haplotype diversity and a lack of fine-scale genetic structure: a case study in the vulnerable estuarine turtle *Malaclemys terrapin*

**DOI:** 10.64898/2026.07.23.740242

**Authors:** Sam Weaver, Tonia S. Schwartz, Iwo P. Gross, Thane Wibbels, Matthew E. Wolak

## Abstract

A central goal when assessing patterns of population structure for conservation is to identify populations with unique genetic compositions. The use of genomic sequencing to identify distinct populations has become an increasingly popular method of delineating conservation units. Reduced costs associated with sequencing make it possible to generate larger, more informative datasets to assess genetic diversity within and among populations. In species that exhibit nest-site philopatry, genetic population structure can emerge on much finer scales, particularly in maternally inherited mitochondrial genomes. We demonstrate the feasibility and insight gained by using whole mitochondrial genome sequencing for evaluating population genetic structure and comparing to previous single marker studies in a vulnerable turtle. We used whole mitochondrial genome sequences from diamond-backed terrapin (*Malaclemys terrapin)* to evaluate whether nest-site philopatry generates fine-scale genetic structure among *M. terrapin* nesting beaches in western Mobile Bay (Alabama, USA). We then compared haplotype diversity between the Alabama population and *M. terrapin* populations from the Atlantic and Gulf coasts and evaluated the utility of using whole mitochondrial genomes rather than a subset of loci to characterize unique haplotypic diversity. We found no genetic structure associated with nest-site philopatry within Alabama, but none of the haplotypes in this region were shared with other Gulf Coast sites. This genetic structure is consistent with strong female natal philopatry within western Mobile Bay relative to the Gulf of Mexico and suggests that the Mobile Bay population is genetically unique relative to other *M. terrapin* populations and merits a unique conservation and management plan.

## Introduction

A central goal of conservation biology is the delimitation and preservation of genetically unique populations across the range of widespread species (Ouborg et al. 2010; Kling et al. 2018), and particularly for those experiencing range-wide declines in abundance (Moritz 1994; Coates et al. 2018; Milot et al. 2020). When populations of widespread species exhibit marked differentiation from geographically proximate populations, they are often given special priority as conservation units (Jensen et al. 2014; Bradshaw et al. 2018; Menchaca et al. 2020). Such conservation units often represent lineages with independent evolutionary trajectories, and management actions to define the geographic scale of genetic diversity and preserve such diversity are important for maintaining evolutionary potential (Moritz 1994; Holderegger et al. 2019). Thus, quantifying genetic diversity and determining the role that dispersal may play in generating observed patterns of population structure is necessary to ensure effective monitoring and conservation of threatened populations (Holderegger et al. 2019; Li and Kokko 2019; (Cavedon et al. 2022).

Diamond-backed terrapins (*Malaclemys terrapin*) are turtles endemic to estuaries along the Atlantic and Gulf coasts of the United States and Bermuda and are classified as vulnerable on the IUCN Red List (Roosenburg et al. 2019). Diamond-backed terrapin populations have been decimated by anthropogenic activities throughout their extensive range, including the historical harvest and shipment of turtles to make terrapin stew in the late 1800s and more recently by nesting habitat loss due to coastal development, drowning as bycatch in traps of commercial crab fisheries, and being struck by vehicles (Converse et al. 2017; Roosenburg and Kennedy 2018; Roosenburg et al. 2019). Like many other long-lived species with well documented threats to long-term persistence, recent conservation strategies have used long-term monitoring and analyses of small sets of markers (e.g., mitochondrial DNA sequence regions and nuclear microsatellites, Avise et al. 1987; Moritz 1994) to quantify population declines, define management units, and estimate key demographic parameters.

Previous conservation genetic studies in diamond-backed terrapins have typically used nuclear microsatellite markers to explore patterns of genetic diversity and patterns of differentiation among populations across the range (Hauswaldt and Glenn 2005; Drabeck et al. 2014; Hart et al. 2014; Petre et al. 2015). These microsatellite data show little evidence for genetic differentiation among populations at small scales (<140 km) (Converse and Kuchta 2018). Past studies using mitochondrial DNA sequences (e.g. ND4 and ND5) to assess patterns of diversity and differentiation among *M. terrapin* populations show clear differentiation between Atlantic Coast populations and those in Texas and Louisiana with little differentiation among populations within those two clusters (Lamb and Avise 1992; Parham et al. 2008). However, these previous genetic studies omitted key parts of the species range, particularly the region from Louisiana to Florida in the Gulf of Mexico. Additionally, these studies are based on short sequence data and marker-based approaches that have known limitations that can be improved upon by analyzing full mitochondrial genomes, which are becoming increasingly inexpensive to sequence and far less expensive than sequencing full nuclear genomes (Aylward et al. 2022; Morón-López et al. 2022). The use of full mitochondrial genomes can also provide deeper insight into population trends, genetic structure, and functional genetic diversity as compared to marker-based analyses (Ouborg et al. 2010; Allendorf et al. 2010).

Here, we explore patterns of genetic structure and variation among nesting beaches of diamond-backed terrapin in western Mobile Bay, Alabama using 19 whole mitochondrial genome sequences. These samples represent a part of the species range not included in previous studies. We quantify mitochondrial haplotype and nucleotide diversity, test for fine-scale (<4 km) spatial genetic structure associated with nesting beaches within Mobile Bay, and assess evidence for past genetic bottlenecks. We then use previously published sequence data from across the species range to assess the degree to which the western Mobile Bay population demonstrates unique genetic diversity relative to the rest of the species range.

## Methods

### Sampling and whole genome sequencing

We obtained blood samples from sexually mature *M. terrapin* females captured along the Mississippi coastline and at multiple beaches along the western coast of Alabama (**Table S1**, **Fig. 1**). Females were discovered during nesting excursions at Cedar Point (Mobile Co., AL) and around Dauphin Island (Mobile Co., AL) during peak nesting season (May-August) using drift fence arrays with pitfall traps or periodic visual encounter surveys. We obtained samples from two female *M. terrapin* inadvertently captured in commercial crab pots in the vicinity of Ocean Springs (Jackson Co., MS). We include these two females in our study for comparison despite not knowing their exact capture location. Considering female diamond-backed terrapins have been documented traveling up to 50km, it is possible they were captured far from their nesting location (Lamont et al. 2021).

**Figure 1:**
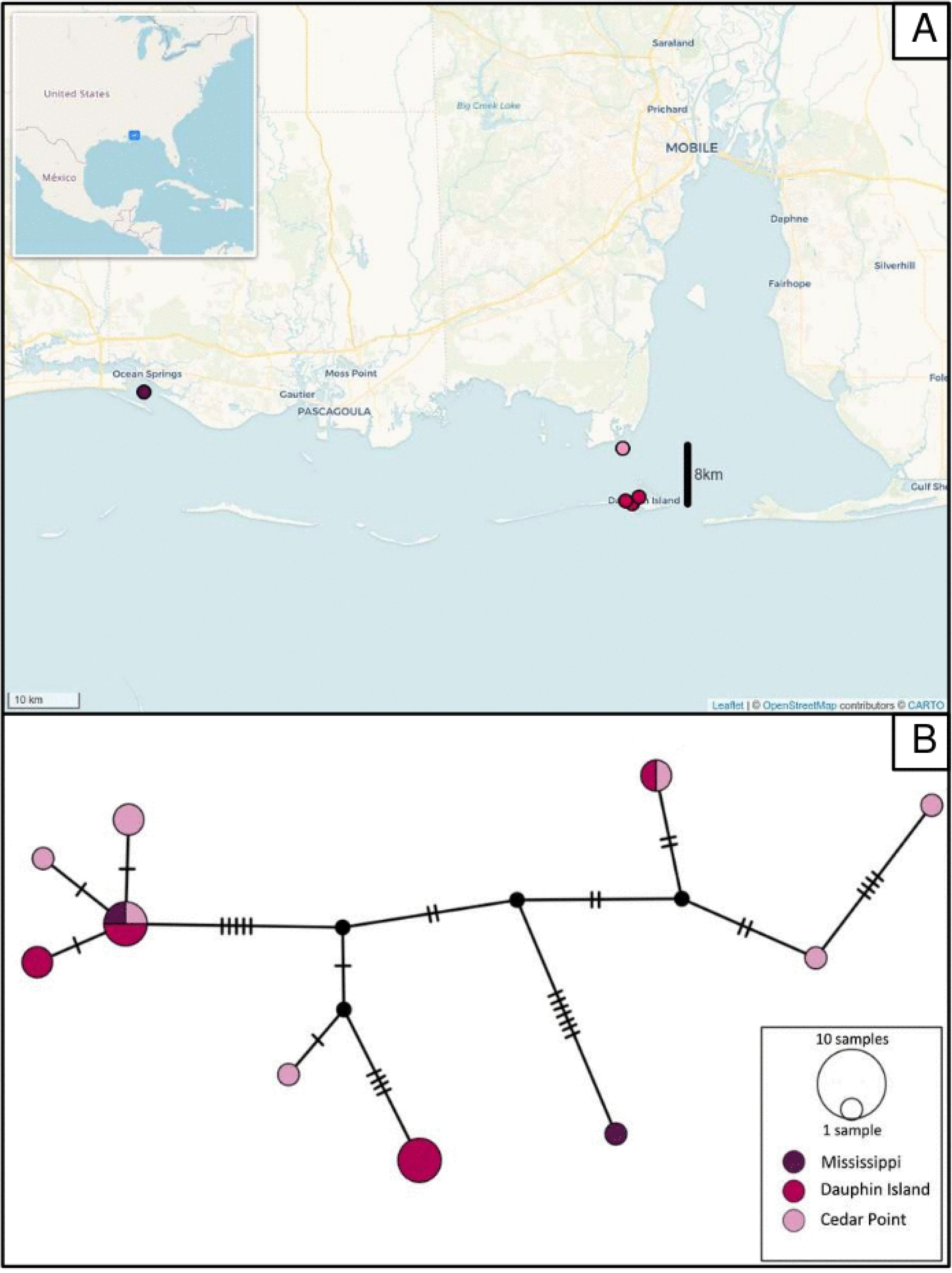
Malaclemys terrapin mitochondrial genetic diversity in western Alabama. **A)** Sampling locations for all 19 Alabama and Mississippi samples color coded by nesting beach complex. Map data from OpenStreetMap under ODbL license https://www.openstreetmap.org/copyright. **B)** Mitochondrial haplotype network representing the number of nucleotide differences (tick marks) between different haplotypes. Haplotypes are colored according to their nesting beach complex(es) (Alabama samples) or location found (Mississippi samples), and circle size represents the number of samples with that haplotype.

Blood samples were obtained from each female diamond-backed terrapin via the subcarapacial venous sinus (Mans 2008), aliquoted into microcentrifuge tubes, and kept on ice in the field until transfer to a −80°C freezer or centrifugation and separation to plasma and red blood cells before freezing. All turtle capture and blood sampling was performed under approved Institutional Animal Care and Use Committee protocol 2019-3539 and Alabama Department of Conservation and Natural Resources collection permits (license numbers 2019066501868680, 2019066503068680, 2020130865668680, 2020130868068680).

DNA was isolated from 15 µl of whole blood or 7.5 µl of a red blood cell pellet using the DNAeasy Kit (Qiagen). We sent 19 samples to Beijing Genome Institute (BGI) for library preparation and whole genome sequencing on the DNBseq platform with 150bp PE reads: 4 samples at 40X coverage and 15 samples at 20X coverage.

### Bioinformatics

Raw reads were processed at BGI in the following way using SOAPnuke (Chen et al. 2018) and software filter parameters “-n 0.001 -| 10 -q 0.4 –adaMR 0.25 --ada_trim” to: trim adaptor sequences allowing two mismatches between the adaptor sequence and raw sequences when identifying adaptors for trimming, low quality reads that contained 40.0% or more of bases with phred scores below 10 were removed from the dataset, and all cleaned reads were rewritten and the read quality value system set to Phred+33.

We obtained the reference sequence for a Maryland *M. terrapin* mitochondrial genome from NCBI (project number PRJNA339452, accession KX774423.1). All reads were aligned to this mitochondrial reference genome using the bwa mem v0.7.17 algorithm (Li 2013). We then used bcftools v1.17 (Li 2011) to call variants from the aligned reads and generate both variant and invariant VCF files and the consensus mitochondrial genomes sequence from each individual.

### Local and rangewide mitochondrial haplotype diversity

For our 19 local samples from Alabama and Mississippi, we generated a Median Joining Haplotype Network in PopArt v1.7 (Bandelt et al. 1999; Leigh and Bryant 2015) to assess the diversity of whole mitochondrial genome haplotypes. We also downloaded a dataset from NCBI (PopSet 171474193) containing 27 individuals from across most of the *M. terrapin* range (**Fig. 2**), but with a notable gap in sampling from Louisiana to Florida (Parham et al 2008). This dataset covers a 2325 bp region of the mitochondrial genome spanning the ND3 and ND4 genes. We extracted this region from our 19 samples and the Maryland reference and created a 2325 bp alignment across all samples using MUSCLE in Geneious v2022.0.2 (https://www.geneious.com). We used this alignment to create a second Median Joining Haplotype Network to visualize the relationships among ND3-ND4 haplotypes and populations using PopArt.

**Figure 2.**
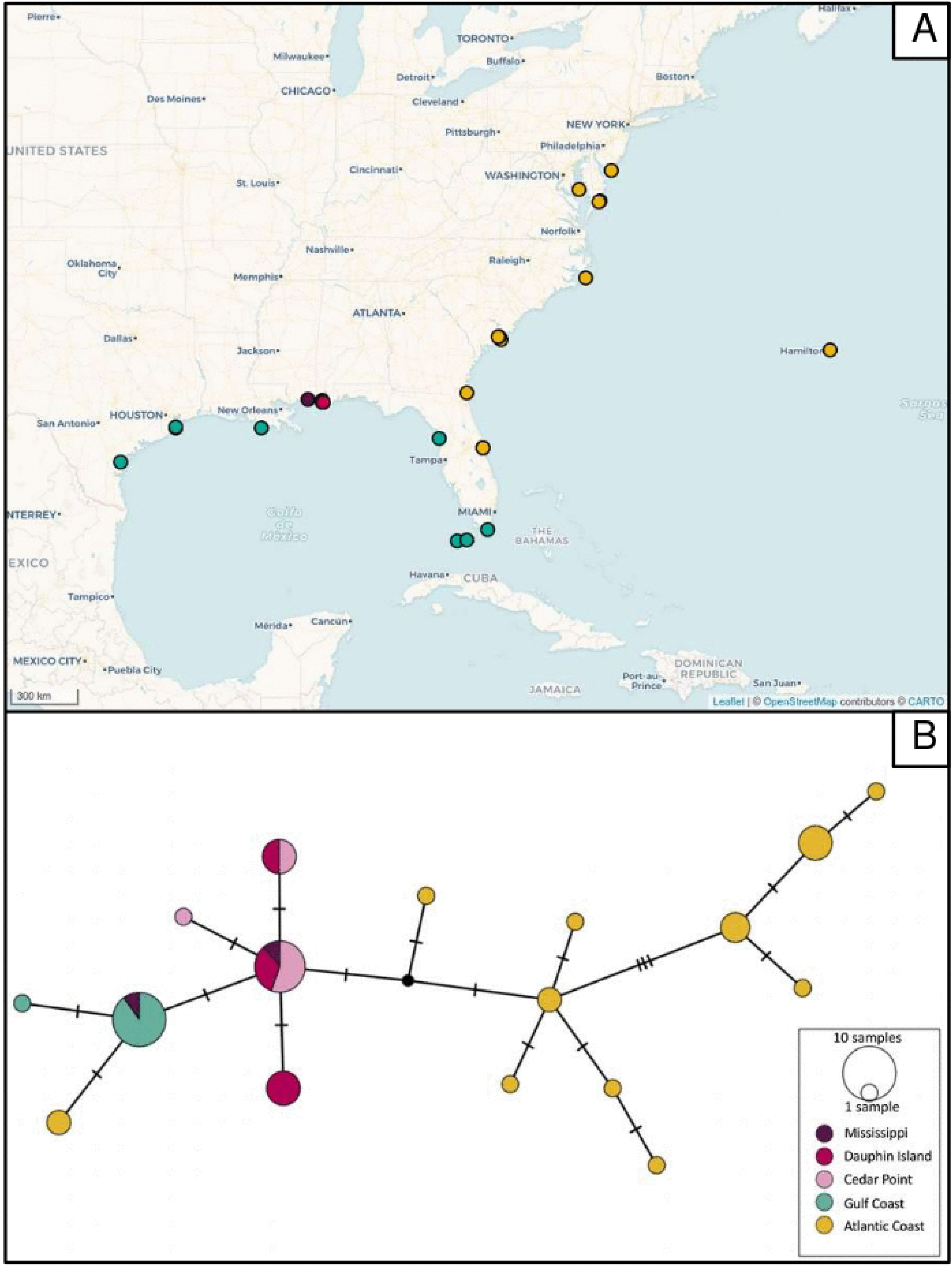
Mitochondrial genetic diversity across the *Malaclemys terrapin* range. **A)** Geographic locations of samples covering genes ND3 and ND4. Map data from OpenStreetMap under ODbL license https://www.openstreetmap.org/copyright. **B)** Mitochondrial haplotype network representing the number of nucleotide differences (tick marks) between different haplotypes. Haplotypes are colored according to their geographic location, and circle size represents the number of samples with that haplotype.

### Population genetics

To assess the degree of genetic structure and differentiation among nesting beaches, we estimated metrics of divergence (D_XY_) and differentiation [Hudson’s F_ST_ (Hudson et al. 1992)] among western Mobile Bay nesting beaches using the PopGenome package (v2.7.5 Pfeifer et al. 2014) in R (R Core Team, 2022). To assess the amount of genetic variation present in the 19 samples, we estimated nucleotide diversity (pi) for each beach by separating individuals into three geographic localities [Dauphin Island, AL (n=9); Cedar Point, AL (n=8); and Ocean Springs, MS (n=2)]. We also estimated pi when all individuals were treated as a single population. We estimated Tajima’s D while treating all samples as one population to see if we could detect signatures of a non-neutral evolutionary history associated with a recent bottleneck followed by population expansion. We used the software MS (October 17 2007 version; Hudson 2002) to simulate 5,000 replicates of expected genotypes under a neutral evolutionary history and compared simulated estimates of Tajima’s D to our observed value to assess whether our observed statistic differed from the simulated values representing a null hypothesis of only neutral evolution.

## Results

We found nine unique mitochondrial genome haplotypes among our 17 Alabama samples, which formed three main haplogroups (**Fig. 1**). Despite the presence of multiple mitochondrial haplotype clusters, there appears to be no genetic structure associated with the geographic distribution of these haplotypes among nesting beaches within western Mobile Bay. This is reflected in our estimates of divergence and differentiation between Cedar Point and Dauphin Island, with low D_XY_ (0.000589) and F_ST_ (0.0227).

Given the lack of evidence for differentiation among beaches in western Mobile Bay, we treated all Alabama samples as one population for characterization of nucleotide diversity and Tajima’s D across the mitochondrial genome. Estimated pi for all Alabama samples was 0.000543, and the observed Tajima’s D for western Mobile Bay samples was slightly negative (−0.294) but well within the range of simulated values for our samples under a neutral evolutionary history (mean = −0.0860, SD = 0.897).

The nine Alabama haplotypes, based on mitochondrial whole genome sequences, condensed to four haplotypes at the ND3-ND4 region, none of which have been observed elsewhere in the species range (**Fig. 2**). One of the Mississippi samples shared the most common Alabama haplotype, whereas the other Mississippi sample clustered with the Texas/Louisiana/Florida Keys haplotypes (Haplotype A in Parham et al 2008). Notably, the Florida central Atantic coast samples (Haplotype D in Parham et al 2008) located at the far left of our haplotype network (**Fig. 2B**), continue to cluster more closely to the other Gulf Coast samples from Parham et al than they do to other Atlantic coast samples from further north. Importantly, in the context of characterizing potentially functional genetic diversity that may contribute to species adaptive potential, from the 75 sequence variants in our whole mitochondrial genome alignment we identified eight non-synonymous SNPs. Four of these show variation within our western Mobile Bay samples and four were fixed alternate alleles relative to the reference mitochondrial genome from an Atlantic coast (Maryland) individual. Only one non-synonymous SNP was found within the ND3-ND4 region (**Table S2**).

## Discussion

Using whole mitochondrial genome sequences, we assessed the genetic status of diamond-backed terrapin populations in western Mobile Bay, Alabama. We found little evidence for genetic structure among nesting beaches. While a relatively small sample size could mask weak structure, the consistent inclusion of individuals from across all nesting beach locations within haplogroups suggests that limited power is not the primary driver of our findings. While species with male-biased dispersal often show strong signatures of population structure in mitochondrial DNA at small geographic scales (Lyrholm et al. 1999; Nishizawa et al. 2011; Converse and Kuchta 2018; Roycroft et al. 2019; Moore et al. 2020), our findings do not point towards genetic evidence for nest-site philopatry in Mobile Bay despite over 6 km of mostly open water between the major nesting areas of Cedar Point and Dauphin Island. This lack of fine-scale structure may also be due to the collection and import of turtles from greater Mississippi, Alabama, and Louisiana into Cedar Point as part of a turtle farming operation in the late 1800s (The New York Times 1881). This recent, widespread admixture could obscure any signature of historical differentiation across beaches within western Mobile Bay resulting in the observed panmixia among our nesting beaches.

Estimated nucleotide diversity within the western Mobile Bay population indicates a surprising lack of genetic diversity, with much lower observed pi (0.000543) than was observed for mitochondrial genomes in populations of the critically endangered Kemp’s Ridley sea turtle (0.00133) (Frandsen et al. 2020). This lack of genetic diversity may be cause for concern in this population, since it potentially limits the capacity of the western Mobile Bay population to persist in the face of threats such as climate change, disease, and the accumulation of deleterious variants (Allendorf et al. 2010; Kosch et al. 2019; Razgour et al. 2019). The observed negative Tajima’s D (−0.294) is consistent with our hypothesis of a past bottleneck due to anthropogenic activity, but this conclusion is not well supported since the value is well within the range of simulated values for our samples under a neutral evolutionary history. Future work incorporating nuclear data with demographic modeling could provide a general model for how ecological drivers and anthropogenic impacts can affect the genetic structure and demographic history of natural populations.

We also sought to clarify the genetic status of the Alabama population in the larger context of the region and the species using a smaller region of the mitochondrial genome. The samples from western Mobile Bay are, on average, less differentiated from other Gulf Coast populations than they are from the Atlantic Coast populations based on our analysis with only the ND3-ND4 region. Even with this smaller region, we find that the western Mobile Bay beaches share no haplotypes with samples from any other location along the Gulf or Atlantic Coasts in the published dataset that we used. Additionally, the use of full mitochondrial genomes more than doubled the number of haplotypes that we observed when using only the ND3-ND4 region and allowed us to identify eight non-synonymous SNPs. Of these eight non-synonymous SNPs, seven were found outside the ND3-ND4 loci and would not have been detectable using the reduced genetic dataset. This additional data would be useful in assessing patterns of nucleotide, amino acid, and haplotypic diversity among populations if applied to a greater number of beaches and sampling localities.

Since we did not detect any of the 15 previously identified mitochondrial haplotypes from Parham et al. (2008) in western Mobile Bay, these findings could be explained as genetic evidence for female philopatry to this previously unsampled area of the Gulf of Mexico, selection against the unique western Mobile Bay haplotypes in other populations, or simply chance differences arising from small sample sizes. Combining the observation that none of the 15 haplotypes from Parham et al. (2008) were found among the western Mobile Bay samples and the fact that our nine whole mitochondrial genome sequence haplotypes reduced down to four haplotypes at the ND3-ND4 region, suggests the range-wide haplotype network (**Fig. 2**) is not capturing the full haplotype diversity at the mitochondrial genome-wide scale. We cannot evaluate the alternative hypotheses explaining the uniqueness of ND3-ND4 region haplotypes in western Mobile Bay, but suggest a range-wide evaluation of mitochondrial genome-wide haplotype diversity is necessary to characterize the extent of female dispersal. While we did not observe any previously identified haplotypes when comparing western Mobile Bay to populations range-wide, female philopatry through movement within western Mobile Bay for selecting nest sites on land was not supported. This further suggests that we have not yet captured the full mitochondrial haplotype diversity in this region of the Gulf of Mexico. Our use of full mitochondrial genomes allowed us to recover more than twice as many haplotypes as would have been identified using a reduced set of loci, and this approach revealed functional diversity that may contribute to adaptive potential of the population and species. It also allowed us to show that nucleotide diversity in western Mobile Bay was markedly lower than observed diversity in other critically endangered species. The uniqueness of haplotypes and poor levels of diversity within this Alabama population suggests this region is of particular concern for conservation and may merit recognition as a distinct conservation unit.

## Acknowledgements

WM Roosenburg graciously provided the biological samples for the Maryland mitochondrial reference genome. We would like to acknowledge M Pop and MP Cummings who generously made the mitochondrial reference genome sequence available on NCBI.

## Data Availability

The raw sequence data files for the 19 individuals are available on NCBI SRA, BioProject PRJNA1483480 with accession numbers listed in Table S1. Alignments and code and input files used to run analyses are available at the Zenodo DOI 10.5281/zenodo.21419405.

## Author contributions

**SW** Formal Analyses, Writing (original draft, review, and editing), Visualization; **TSS** Investigation, Resources, Formal analyses, Writing (review and editing), Project administration; **IPG** Resources, Writing (review and editing); **TW** Resources, Writing (review and editing); **MEW** Conceptualization, Methodology, Resources, Writing (review and editing), Project administration, Funding acquisition, Visualization.

## Supplemental Captions

**Table S1:** Individual *Malaclemys terrapin* sample identities and locations for all individuals sequenced from the Alabama and Mississippi populations. Statistics from mapping sequence read data to the reference mitochondrial genome for each individual calculated using QualiMap (Okonechnikov et al. 2025). The Accession column contains the National Center for Biotechnology Information Sequence Read Archive (NCBI SRA) accession numbers from BioProject PRJNA1483480 of the read data used in these analyses. Sample location and voucher information for the NCBI data from 27 individuals’ mitochondrial ND3 to ND4 region can be found in the Electronic Supplementary Materials to Parham et al. (2008).

**Table S2:** Characteristics of sequence variants found in the protein-coding genes across the whole mitochondrial genome sequence alignment (19 Alabama and Mississippi samples) relative to the Maryland reference sequence. The Type column refers to the type of sequence variant using the following abbreviations: single nucleotide variant (SNP), transition (TS), transversion (TV). The Frequency column refers to the frequency of the Gulf of Mexico individuals in our sample that contained the sequence variant. The Pattern column refers to whether the sequence variant was found in some frequency within our 19 Alabama and Mississippi samples, or fixed in all the Alabama and Mississippi samples relative to the Maryland reference genome.

## Notes

### Competing Interest Statement

The authors have declared no competing interest.

